# State-dependent changes in olfactory cortical networks via cholinergic modulation

**DOI:** 10.1101/452003

**Authors:** Donald A. Wilson, Maxime Juventin, Maria Ilina, Alessandro Pizzo, Catia Teixeira

**Author notes:** **Address for Correspondence:** Donald Wilson Emotional Brain Institute Nathan Kline Institute for Psychiatric Research 140 Old Orangeburg Road Orangeburg, NY 10962.

## Abstract

Activity in sensory cortical networks reflects both peripheral sensory input and intra‐ and inter-cortical network input. How sensory cortices balance these diverse inputs to provide relatively stable, accurate representations of the external world is not well understood. Furthermore, neuromodulation could alter the balance of these inputs in a state‐ and behavior-dependent manner. Here, we used optogenetic stimulation to directly assay the relative strength of bottom-up (olfactory bulb) and top-down (lateral entorhinal cortex) synaptic inputs to piriform cortex in freely moving rats. Optotrodes in the piriform cortex were used to test the relative strength of these two inputs, in separate animals, with extracellular, monosynaptic evoked potentials. The results suggest a rapid state-dependent shift in the balance of bottom-up and top-down inputs to PCX, with enhancement in the strength of lateral entorhinal cortex synaptic input and stability or depression of olfactory bulb synaptic input during slow-wave sleep compared to waking. The shift is in part due to a state-dependent change in cholinergic tone as assessed with fiber photometry of GCaMP6 fluorescence in basal forebrain ChAT+ neurons, and blockade of the state-dependent synaptic shift with cholinergic muscarinic receptor activation.

---

Activity in sensory cortical networks reflects both peripheral sensory input and inputs from intra‐ and inter-cortical connections ^1-6^. These intra/inter-cortical networks convey information about sensory context, expectation and internal state, which can shape cortical response to sensory input ^4,7-10^. As has been demonstrated in essentially all sensory systems, this means that often very early in the sensory processing stream context, internal state, and past experience can shape sensory coding. Similar state‐ and experience-dependent modulation occurs in many systems ^11-15^, but precisely how the sensory cortex balances these diverse inputs to provide relatively stable, accurate representations of the external world is not well understood.

In primary olfactory (piriform; PCX) cortex these sensory and intra/inter-cortical inputs, which are primarily glutamatergic, are strictly segregated into distinct lamina, with sensory input from the olfactory bulb targeting the Layer II/III pyramidal cell distal apical dendrites in Layer Ia, and intra/inter-cortical inputs targeting the proximal apical dendrites in Layer Ib and basal dendrites in Layer III ^16^. Both inputs also target several classes of inhibitory interneurons ^17^. The synapses in these two lamina differ dramatically. For example, Layer Ia excitatory synapses are relatively much stronger than Ib ^18^, yet show substantially less activity-dependent long-term synaptic potentiation ^19,20^. Furthermore, Layer Ib (and presumably layer III) synapses are under significantly greater control of neuromodulators such as acetylcholine (ACh) and norepinephrine (NE) than their counterpart synapses in Layer Ia ^21-23^. For example, activation of pre-synaptic ACh muscarinic receptors on Layer Ib fibers induces synaptic depression, but has little or no effect on pre-synaptic release from Layer Ia fibers ^21^. In addition, Layer Ib but not Layer Ia synapses express pre-synaptic GABA_B_ receptors, which allows GABAergic presynaptic modulation of intra‐ and inter-cortical inputs to PCX but not of olfactory bulb inputs ^24,25^. Together, these findings suggest a differential regulation of Layer Ia and Ib/III inputs based on context, internal state and past experience.

In accord with the concept of sensory afferent (Ia) – top down (Ib/III) balance, indirect functional connectivity assays, such as local field potential coherence ^26^ and functional MRI ^27^ suggest a shift in net connectivity from the olfactory bulb to neocortical structures as rodents shift from waking/fast-wave state to sleeping/slow-wave state. It has been hypothesized that this shift may be critical for sleep-dependent odor memory consolidation. In fact perfusion of the PCX with the GABA_B_ agonist baclofen during post-training sleep and memory consolidation, which depresses layer Ib synapses ^24,25^, impairs accurate odor memory ^25^.

Here, we more precisely explore the state-dependent shift in PCX layer Ia and Ib/III balance by using a direct assay of synaptic connectivity. Specifically, we used optogenetic activation of identified inputs to layers Ia (i.e., olfactory bulb CAMKII-expressing mitral/tufted cells) and Ib/III (i.e., CAMKII-expressing lateral entorhinal cortical [LEC] neurons) within the PCX in awake, freely moving rats as they naturally cycled between sleep and waking states as assessed with local field potentials. The optical stimulation allowed induction of monosynaptic extracellular evoked potentials that varied in strength in a state‐ and ACh-dependent manner. The results suggest a rapid and robust state-dependent shift in the balance of bottom-up and top-down synaptic inputs to PCX.

## Results

An example of the laminar separation of OB and top-down inputs to PCX is shown in Fig.1 where CAMKII-YFP virus labels OB inputs to the PCX and CAMKII-mCherry labels top-down inputs to the PCX. All other rats had virus in either OB or LEC, not both. Data from a total of 6 rats with AAV-ChR2 transfections targeting CAMKII expressing cells in the LEC and 9 different rats with transfections targeting CAMKII expressing cells in the main OB (4 chronic recordings and 5 urethane anesthetized recordings) are included here. In both cases, optotrodes were implanted in the PCX to record either top-down (LEC->PCX) or bottom-up (OB->PCX) extracellular, light-evoked synaptic potentials (EP’s). Light-evoked PCX EP’s were recorded in either the rat’s home cage or in a Plexiglas circular arena lined with standard bedding to which the animal had been habituated. Recording sessions lasted 1-4 hrs. primarily during the light phase of the light-dark cycle, during which time the animals naturally cycled between sleep-wake states. Light-evoked EP’s were triggered in the PCX at 0.05Hz throughout the recording session. This frequency of stimulation did not induce any detectable decrease in response amplitude over the course of the session (within a specific state).

**Figure 1.**
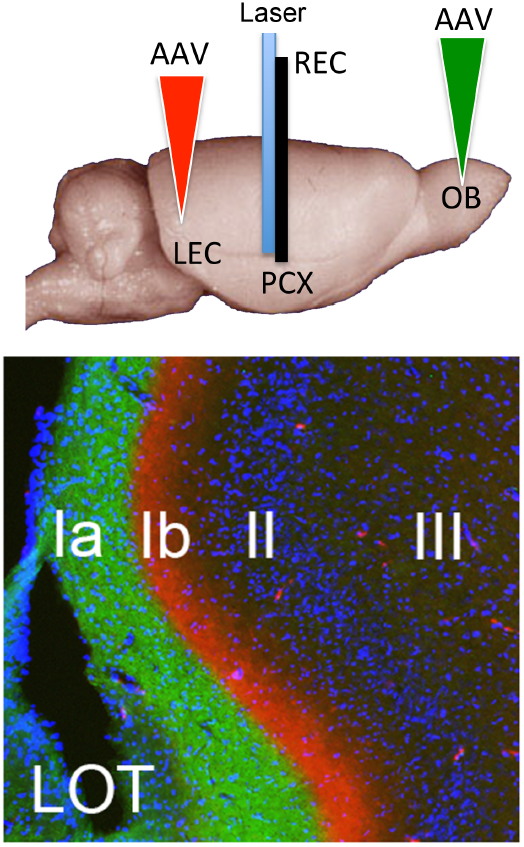
(**TOP**) Animals used for data collection had AAV-ChR virus injected into either the OB or the LEC, and an optotrode implanted in the PCX to record extracellular potentials evoked by light stimulation of either fibers originating in the OB or, in separate animals, in the LEC. (**BOTTOM**) Example of lamination of bottom-up (green YFP from OB infection) and top-down (red mCherry infection of cortical sites) to the PCX. In individual animals used for data collection, either the OB or LEC were transfected with ChR-eYFP, not both.

A representative LEC infection is shown in Fig. 2A. Cells throughout all layers and their processes were labeled. Axons from these cells projected, among other places, to the PCX where they predominantly targeted layer Ib and to a lesser extent layer III. A 1 ms flash within PCX evoked both unit firing (Fig. 2B) and an extracellular EP in PCX. The EP could reverse polarity from positive to negative as the optotrode moved from layer III to I, respectively, as is similar to electrically evoked potentials ^28^. Given the size of our electrodes, the majority of recordings did not allow isolation of unit activity, and thus no quantification was attempted. As shown in Fig. 2 C and D, the efficacy of LEC->PCX EP’s in the PCX varied with behavioral state. LEC fiber EP’s in the PCX increased in amplitude during slow-wave sleep (i.e., periods of LFP high delta power) compared to fast-wave states (i.e., periods of LFP low delta power) such as waking. Responses during REM appeared similar to those during waking (Fig. 2D), but the brevity and rarity of REM bouts precluded quantitative analysis of REM sleep effects here. There was a significant, mean positive correlation between LEC->PCX EP amplitude and LFP delta band r.m.s amplitude across animals, with 6/6 animals showing individual positive correlations (individual r values converted to z-scores, significantly different from 0, t(5) = 2.52, p = 0.026).

**Figure 2.**
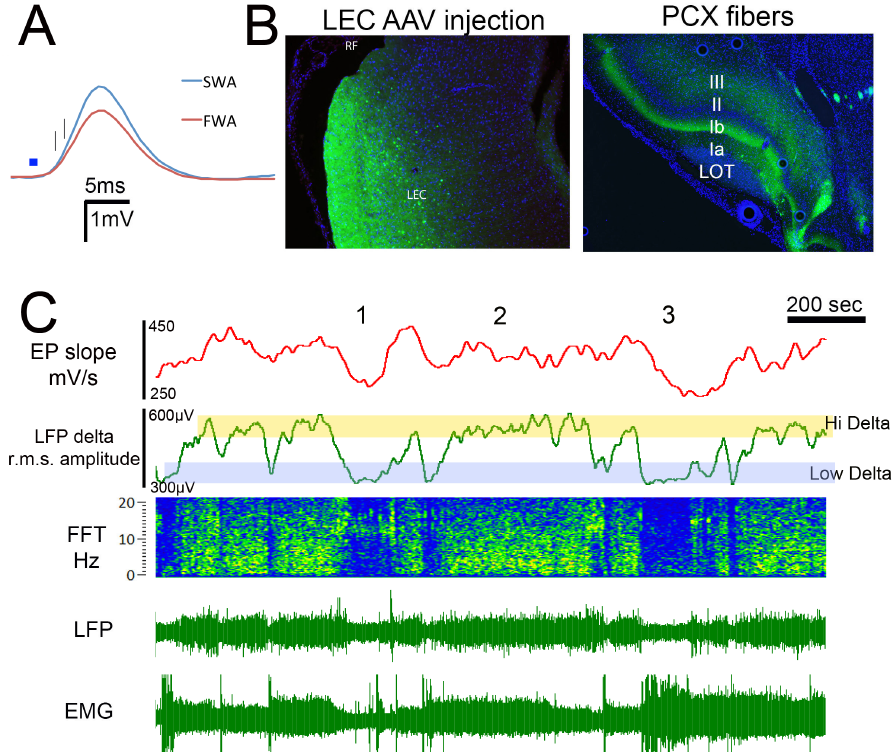
Representative example of state-dependent modulation top-down (LEC->PCX) synaptic efficacy in the PCX. (**A**) Site of AAV-CAMKII-ChR LEC infection and YFP labeled fibers within the PCX. Labeled cell bodies were generally located throughout all layers in LEC, with labeled dendritic and axonal fibers similarly broadly distributed. RF signifies the rhinal fissure. For physiological recordings, the optotrode was placed in PCX to selectively activate axons originating in the LEC. (**B**) Example of light-evoked extracellular EP recorded in PCX layer I/II in response to activation of LEC pre-synaptic fibers (blue square) and example of simultaneously light-evoked unit activity (raster plot and histogram). Given the size of our electrodes, the majority of our recordings did not allow isolation of PCX unit activity. The delay and variable latency of evoked spikes is consistent with a post-synaptic response. (**C**) EP amplitude was determined by using slope measurements (vertical lines) on the early monosynaptic phase of the EP. EP slope varied in a state-dependent manner. (**D**) Recording of PCX local field potential (LFP) and neck muscle EMG in this animal that allowed sleep-wake cycle staging. Delta band power and EMG allowed identification of sleep and wake states. Periods of high and low delta are marked by shading. Individual examples of each state, determined from delta amplitude and EMG activity are highlighted: REM = 1, NREM = 2, waking = 3. As delta band r.m.s. amplitude increased during NREM sleep, the slope of light-evoked extracellular synaptic potentials induced by activation of LEC fibers within PCX also increased. During both REM and waking, EP slope decreased.

In a subset of animals, LFP coherence between the PCX and LEC in the beta frequency band was assessed during the same recording session as the light-evoked EP’s. The enhancement in the optogenetic assay of synaptic efficacy during slow-wave sleep bouts compared to waking was associated with an increase in PCX-LEC coherence (n=4, mean coherence during waking = 0.55± 0.13, during slow-wave sleep coherence = 0.63, t(3) = 2.43, p = 0.047), as previously described ^26^. Similar results were observed in the delta band (0-4 Hz). Thus, during slow-wave sleep, there is both an enhancement in coherence of activity in the LEC and PCX, and an enhancement in the efficacy of LEC fiber synapses within the PCX.

In contrast to this top-down pathway, the efficacy of bottom-up OB->PCX EPs in the PCX was stable or even negatively correlated with LFP delta band r.m.s. amplitude within animals (Fig. 3). This pattern was observed in 4/4 chronically recorded animals. AAV-ChR infection appears to primarily target mitral and tufted cells including their dendritic tufts which extended into glomeruli. Axons of these cells targeted layer I1 within the PCX (Fig. 3A). As with LEC AAV-ChR infection 1ms light flashes within PCX could evoke both single-unit firing and extracellular EP (Fig. 3B). Activation of mitral/tufted cell axons in the PCX evoked stronger EP’s during waking compared to during slow-wave sleep (Fig. 3A, C).

**Figure 3.**
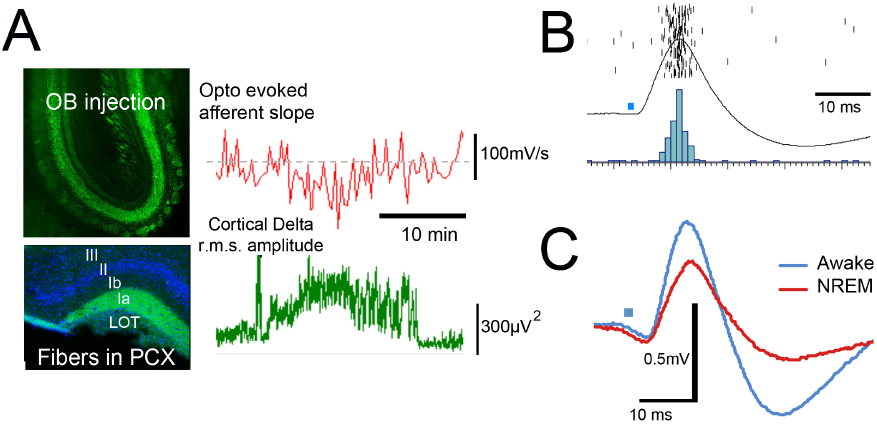
State-dependent modulation of OB inputs to the PCX. (**A**) Example of AAV-CAMKII-ChR infection within OB and YFP labeled axons within PCX. The optotrode was placed within PCX to selectively activate axons originating within the OB. As LFP delta band r.m.s. amplitude increased, indicative of NREM sleep, the slope of light-evoked extracellular synaptic potentials induced by activation of OB fibers in the PCX was stable or decreased. (**B**) Example of evoked potential in PCX in response to light activation (blue square) of mitral/tufted cell axons in PCX. This stimulation evoked both an extracellular potential and unit spiking (rasterplot). (**C**) Examples of light-evoked potentials within PCX during waking and NREM sleep.

Figure 4, shows a direct comparison of state-dependent changes in LEC-> PCX and OB-> PCX synaptic efficacy dependent on delta r.m.s. amplitude in unanesthetized, chronically recorded rats, as well as a group of urethane-anesthetized animals that also cycle between low delta and high delta states ^29,30^. The urethane-anesthetized animals were added since 1) urethane-anesthetized state-dependent changes in PCX odor responsiveness have been extensively examined in the past ^29,30^ and 2) optogenetic responses appeared to show less variation during urethane anesthesia which may allow detection of small differences between states. The OB->PCX chronically recorded rats and the anesthetized rats showed the same state-dependent effects ‐ 2/4 chronically recorded animals and 3/5 anesthetized animals showed an individually significant negative correlation between EP slope and LFP delta amplitude. A one-way ANOVA across correlations (individual r values converted to z-scores) in all three groups revealed a significant effect (F(2,12) = 8.16, p = 0.006). Post-hoc Tukey tests revealed that the LEC->PCX correlation between delta amplitude and EP slope was significantly different from both the chronic and the urethane anesthetized OB->PCX pathways (p < 0.05).

**Figure 4.**
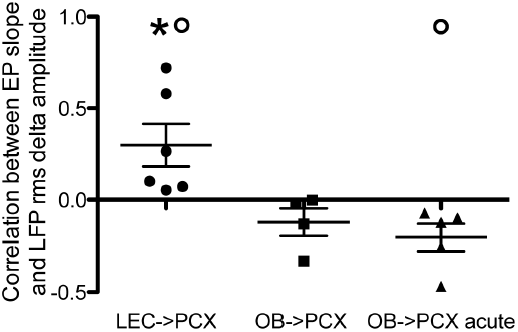
LEC->PCX synaptic input was significantly positively correlated with PCX delta band r.m.s. amplitude, while OB->PCX synaptic input was slightly negatively correlated with PCX delta band r.m.s. amplitude. The OB->PCX depression was also evident in urethaneanesthetized animals that naturally cycle between fast‐ and slow-wave states. Asterisk signifies significant difference between OB and LEC inputs based on one-way ANOVA and post-hoc Tukey tests. Circles represent significant difference between mean correlation and zero. Statistics performed on z-score converted r values.

One potential mechanism of this state-dependent modulation of synaptic input to the PCX is the state-dependent change in activity of basal forebrain cholinergic neurons projecting to the PCX and elsewhere ^23,31,32^. ACh, via muscarinic receptors can pre-synaptically reduce glutamate release from intra‐ and inter-cortical axons in Layer Ib of the PCX ^21^. Thus, during periods of low cholinergic release, such as slow-wave sleep, these intra‐ and inter-cortical fibers should be released from pre-synaptic inhibition.

In order to confirm state-dependent modification of ACh, we used fiber photometry of basal forebrain ChAT+ neurons using GCaMP6 in Long Evans-Tg(ChAT-Cre)5.1Deis rats injected with AAV1.Syn.Flex.GCaMP6s.WPRE.SV40 virus (Fig. 5A). ChAT+ neurons responded to sensory inputs such as sound or odors (sampling 381Hz; Fig. 5B), and showed strong state-dependent modulation (sampling 0.1Hz; Fig. 5C). GCamP6 fluorescence in ChAT+ basal forebrain neurons (with variation in the control channel subtracted to remove potential movement artifacts) showed a negative correlation with LFP delta r.m.s. amplitude (Fig. 5D and E). As LFP delta amplitude increased (NREM state), ChAT+ neuron activity decreased in 3 out of 3 rats (each animal showed a significant r value). The mean of z-score converted correlations was significantly different from 0 (t(2) =6.42, p = 0.023). The mean delta F during waking relative to NREM was 3.1%, a significant increase (Fig. 5F; t(2) = 4.49, p = 0.046). Thus, activity of ChAT+ neurons increases during waking and decreases during NREM, providing a potential mechanism of state-dependent synaptic modulation of inputs to PCX.

**Figure 5.**
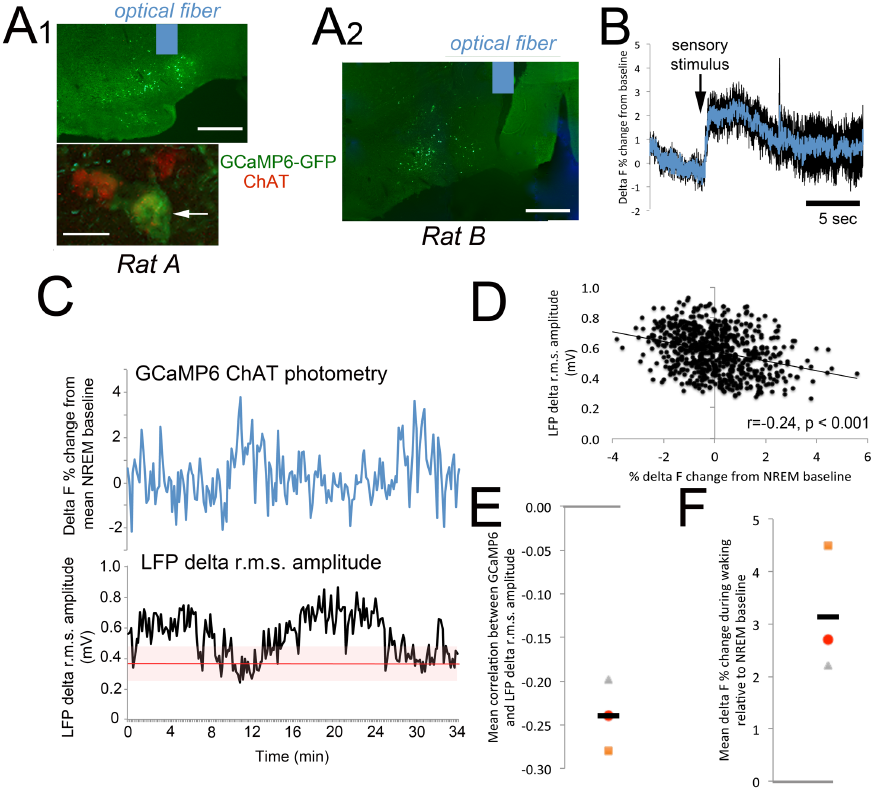
Fiber photometry recording of GCaMP6 fluorescence in ChAT-Cre:GCamp6 freely moving rats cycling between sleep/wake. (**A1**) Top panel: Representative example of GCaMP6-GFP labeled cells in basal forebrain [scale = 500um] and (bottom panel) double labeled GCAMP6-GFP/ChAT immunolabeled cells [arrow, scale=20µm]. Blue rectangle represents approximate location and size of optical fiber used for photometry. (**A2)** Representative example of GFP labeled cells and approximate location of optical fiber in a second animal [scale = 500um]. (**B**) Basal forebrain ChAT+ neurons GCaMP6 response to a mixed auditory/olfactory stimulus in a freely moving (black = SEM). Sampling frequency was 381Hz. (**C**) Example of simultaneous LFP recording and fiber photometry (0.1Hz sampling) monitoring of GCaMP6 fluorescence in basal forebrain ChAT+ neurons in ChAT-cre rats. GCaMP6 fluorescence is expressed as % change from mean NREM levels and fluctuations in the control 405nm channel were subtracted to remove movement artifacts. GCaMP6 fluorescence (i.e., activity of ChAT+ neurons) increased during periods of low delta r.m.s. amplitude (fast-wave or waking state) and decreased during periods of high delta r.m.s. amplitude (NREM). (**D**) Example from one rat showing that GCaMP6 fluorescence in ChAT+ basal forebrain neurons was negatively correlated with LFP delta power; low during NREM and high during waking. (**E**) There was a significant negative correlation between LFP delta r.m.s. amplitude and basal forebrain GCaMP6 ChAT+ neuron fluorescence. All 3 animals showed a similar effect (one sample t-test, p = 0.023). (**F**) Across 3 rats mean GCaMP6 fluorescence (activity of ChAT+ neurons) was higher during waking than the mean activity during NREM sleep which was taken as baseline (one sample t-test, p = 0.046).

To test the role of ACh muscarinic receptors in the state-dependent synaptic changes, a subset (n=6) of the same animals described above were systemically injected with scopolamine (0.5mg/kg) or saline, and again allowed to freely cycle between sleep-wake states for 1-3 hours post-injection. EP data were collected beginning at least 30 min post-injection. As previously reported in humans ^33^, scopolamine significantly enhanced low frequency oscillations, and this enhancement was present during both waking (Fig. 6A; ANOVA frequency X drug interaction, F(41,246) = 4.12, p = 0.03) and NREM sleep (Fig. 6B; F(41,246) = 4.95, p < 0.0001). Sleep-wake cycles appeared normal under scopolamine.

**Figure 6.**
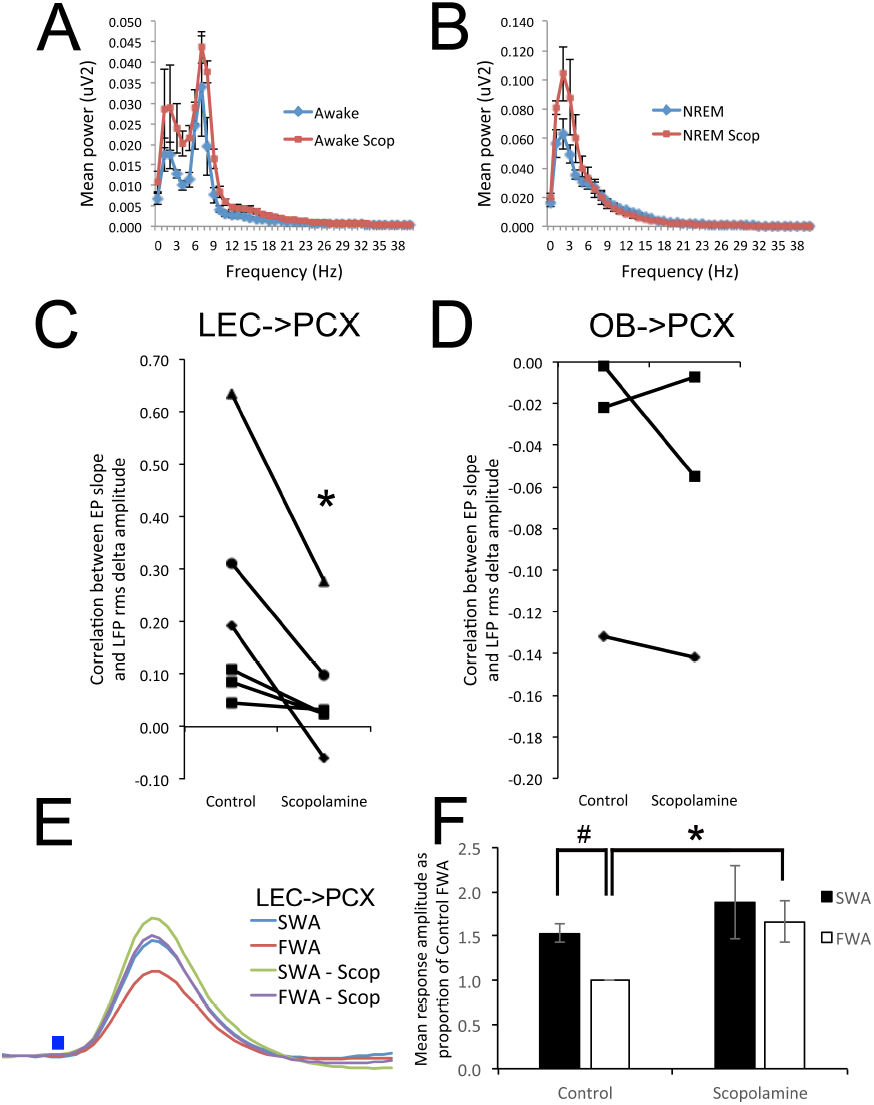
The state-dependent modulation of LEC->PCX synaptic strength was cholinergic dependent. FFT analyses of the effects of scopolamine PCX LFP’s during waking (**A**) and NREM sleep (**B**). During both waking and NREM, scopolamine induced an enhancement in delta power (drug X frequency ANOVA, significant interaction), as previously reported ^33^. (**C**) The muscarinic receptor antagonist scopolamine significantly reduced the dependence of LEC->PCX EP slope on sleep/wake state. 6 of 6 animals showed a reduced correlation between LEC->PCX slope and r.m.s. delta amplitude. Asterisk signifies significant paired t-test, p = 0.022). (**D**) There was no significant effect of scopolamine on OB->PCX evoked response slope (paired t-test, N.S.). (**E**) Examples of light-evoked responses in PCX in response to LEC fiber activation during slow-wave (SWA) and fast-wave (FWA) activity in the same animal before and after scopolamine (SCOP). Both slow-wave sleep and scopolamine induce a response increase. (**F**) Mean EP slopes (as proportion of control FWA) during sleep and waking show a significant difference in control animals (hashtag signifies significant difference between SWA and FWA, ttest, p < 0.05), but no significant difference during scopolamine. Asterisk signifies significant difference in EP slope during FWA between control and scopolamine treated animals.

As shown in Fig. 6C, state-dependent modulation of top-down, LEC->PCX EP’s was significantly attenuated by scopolamine. There was a significant decrease in the positive correlation between EP amplitude and LFP delta r.m.s. amplitude after scopolamine injection (t(5) = 2.68, p = 0.022). This flattening of the state-dependent modulation was primarily due to a scopolamine-induced enhancement in EP amplitude during waking (response during control waking vs. scopolamine waking; t(5) = 2.85, p = 0.018), thus reducing the relative difference between waking and slow-wave sleep EP’s (Fig. 6E, F). Interestingly, no significant effect of scopolamine was detected in the bottom-up, OB->PCX pathway (n=3, control, mean r = −0.12, Scop, mean r = −0.07, t-test N.S., Fig. 6D).

## Discussion

The present results demonstrate a state-dependent shift in the balance between identified bottom-up and top-down projections to primary sensory cortex. During slow-wave sleep-related periods of high amplitude delta oscillations, efficacy of LEC synapses within the PCX is enhanced, in part via a cholinergic-dependent mechanism. During the same state, efficacy of OB synapses within the PCX is unchanged or weakly depressed. The mechanism of this OB modulation is not known, though it corresponds to the decrease in PCX odor-evoked responsiveness observed during slow-wave sleep ^29,34^. Unlike the LEC-PCX modulation, acetylcholine muscarinic receptor blockade does not significantly impact the OB->PCX state-dependent shift. These state-dependent shifts may be critical for sleep-dependent olfactory memory consolidation, allowing odor memory replay to occur within the context of top-down associative activity, in the absence of interference from ongoing bottom-up odor input ^35^. Given the observed role of ACh in these effects, we hypothesize that similar shifts in top-down/bottom-up balance may occur during less pronounced state changes, such as quiet awake versus vigilance, or task engagement versus disengagement, as suggested by indirect measures of functional connectivity ^12,36^. Work on this hypothesis is ongoing.

The optically evoked potentials measured here resemble classic electrically evoked mono-synaptic potentials described in the PCX ^16,37^. Given that both olfactory bulb and top-down inputs can target pyramidal cell dendrites and inhibitory interneurons ^16,17,38,39^, the extracellularly recorded evoked potentials presumably reflect inputs to both cell classes. Whether synapses on both classes of target cells show similar state-dependent modulation is not known, however, cholinergic synaptic modulation in this system can be pre-synaptic ^21^, and thus may modulate both targets. Future work is required to identify whether synapses on different cell types are modulated similarly, and whether the observed state-dependent changes in synaptic strength correspond to changes in evoked post-synaptic spiking. In addition, while optogenetic stimulation could induce antidromic and/or polysynaptic activation, our slope measures were taken at short latencies, prior to the onset of these polysynaptic evoked responses, as is standard for electrically evoked synaptic potential measures. Finally, recent evidence also suggest that there may be an inhibitory projection from the entorhinal cortex, at least targeting the hippocampal formation ^40^. This pathway is presumably not involved in the responses here since the optogenetics targeted CAMKII expressing neurons, however the state-dependent effects on this pathway are unknown.

These data significantly extend previous indirect assays of functional connectivity changes associated with sleep, by directly examining synaptic efficacy within identified pathways. There is an extensive literature on functional connectivity changes with sleep, in both humans and non-human animals models ^13,26,41-49^. However, these indirect measures (e.g., local field potential coherence and fMRI functional connectivity): 1) do not allow isolation of specific pathways; 2) provide measures of net changes in functional connectivity which could include bidirectional variation; and 3) cannot distinguish between correlated changes in overall activity (e.g., firing rate) and synaptic efficacy. In contrast, the optogenetic assay used here allows analysis of synaptic efficacy in specific, directional pathways within specific target structures. Current work is examining other pathways into and out of the PCX in different behavioral states.

The data presented here further emphasize that activity of basal forebrain cholinergic neurons fluctuates across the sleep-wake cycle (Fig. 4; ^50-52^). Cholinergic projections to the PCX play a major role in pyramidal cell excitability ^53^, synaptic plasticity ^31,54^, and odor memory and perception ^23,55^. The data here demonstrate that ACh also plays a critical role in state-dependent synaptic connectivity between the PCX and top-down projections from the LEC. Such top-down input may have multiple effects on PCX activity. For example, LEC lesions or activity suppression can enhance odor-evoked PCX single-unit responses, local field potential oscillations ^56^, and PCX odor-evoked immediate early gene activation ^57^, suggesting a tonic top-down suppressive effect of LEC on PCX. In fact, lesions of the LEC can enhance simple odor memory ^58,59^. However, in contrast to the effects on simple odor recognition, LEC suppression impairs well-learned, fine odor discrimination ^56^, and pharmacological suppression of PCX intra- and inter-cortical synapses, including those from the LEC, impairs the accuracy of post-training odor memory consolidation ^25^. Thus, by modulating the synaptic efficacy of LEC, among other, top-down inputs to the PCX, state-dependent changes in ACh could shift the PCX between states of high simple odor learning (low LEC synaptic input), fine sensory discrimination (moderate LEC synaptic input), and odor memory consolidation (high LEC input during slow-wave sleep) abilities ^23,32^.

Finally, it should be emphasized that the state-dependent changes observed here are just that – state-dependent. For example, the enhancement in LEC->PCX synaptic input during NREM sleep does not appear to be a long-lasting re-set of synaptic strength. Thus, these data are not inconsistent with the synaptic homeostasis hypothesis (SHY) of synaptic downscaling during NREM sleep ^60-62^. However, neuromodulatory changes in the strength of specific synaptic inputs, as shown here, could make subsets of synapses within a given pathway more or less susceptible to such homeostatic or experience-dependent changes ^32^. This could even allow strengthening of some synapses ^63^ and weakening of others ^62^ during the same NREM episode, depending on the specific balance of pre‐ and post-synaptic excitation.

In summary, the results described here demonstrate that PCX networks function in a very dynamic state-dependent context of varying bottom-up and top-down inputs. These state-dependent changes may include variation in regional activity levels, but also, as shown here, in the synaptic strength of inputs from these areas to the PCX. Impairments in these dynamic shifts could contribute to impaired odor perception and/or memory.

## Methods

Long Evans hooded male rats were used as subjects. Animals were housed with *ad lib* food and water and a 12:12 hr light:dark cycle. Testing was performed during the light portion of the cycle. All procedures were approved by the NKI IACUC and were in compliance with NIH guidelines.

### Electrophysiology and optogenetics

For channelrhodopsin transfection, animals were anesthetized with isoflurane and holes drilled over either the olfactory bulb or LEC. A glass pipette was used to infuse 0.5-1.0µL of AAV5-CaMKII-hChR2(E123T/T159C)-eYFP from UNC Vector Core. Post-operative treatment included subcutaneous Buprenorphine-SR and enrofloxacin antibiotic. At least 3 weeks later (typically 4-6 weeks), animals were again anesthetized with isoflurane and implanted with optotrodes (127µ stainless steel electrode attached to 200µ optical fiber) aimed at the piriform cortex layer III under physiological control. Extracellular evoked potentials to 1ms light flashes (5-9mW, 473nm, SLOC Lasers) were monitored as the electrode was lowered to the site of maximal response amplitude. The optotrode was then affixed to the skull with dental cement and attached to a subcutaneous telemetry transmitter (DSI, F20-EET) for recording of the PCX optotrode and either local field potentials in the lateral entorhinal cortex or, in different animals, neck muscle EMG. Data were collected (5kHz) and analyzed using Spike2 software (CED, Inc.).

Following recovery, optotrodes were connected to the laser with an optical cable and optical swivel, and the animal placed either in their home cage or in a circular chamber with wood chip bedding (29cm diameter X 36 cm tall) to which it had previously been habituated. Light flashes (0.05Hz, 1ms duration, 5-9mW) were delivered throughout a 1-4hr recording session and the amplitude (initial slope) of light-evoked potentials quantified. The spontaneous PCX local field potential (LFP) was also used to determine behavioral state as the animal cycled between waking (low delta frequency band [0-4Hz] and high theta band [5-15Hz] power) and slow-wave/non-rapid eye movement (NREM) sleep (high delta frequency band power) ^34^. In some animals, EMG was used to identify REM bouts, though these were generally too short and infrequent to allow thorough analyses across animals with the stimulation paradigm used here.

In a subset of animals with olfactory bulb transfections, rather than being prepared for chronic recordings, the animals were anaesthetized with urethane (1.25g/kg), optotrodes implanted into the PCX and light-evoked responses obtained at 0.05Hz as described for the chronic recordings. Animals under urethane anesthesia display fast-wave/slow-wave cycling similar to waking NREM sleep cycles ^29,30^. The relationship between light-evoked response slope and delta power was analyzed as for the unanesthetized animals.

Given the focus here on differences between waking and slow-wave sleep, modulation of evoked potentials by behavioral state was determined by correlating response slope with LFP delta r.m.s amplitude (0.05 Hz samples). Correlations were converted to z-scores for statistical analyses. In addition, in a subset of animals state-dependent changes in LFP coherence between the PCX and the LEC was calculated using the cohere script in Spike2, to allow comparison of identified synaptic change with LFP coherence.

### Fiber photometry

We used fiber photometry to quantify state-dependent changes in activity of ChAT+ neurons in the basal forebrain/horizontal limb of the diagonal band of transgenic rats (Long Evans-Tg(ChAT-Cre)5.1Deis; Rat Resource and Research Center, Columbia Missouri). Under isoflorane anesthesia, ChAT-cre rats (n=3) had AAV1.Syn.Flex.GCaMP6s.WPRE.SV40 virus (2µ) infused in the HLDB basal forebrain to infect ChAT+ neurons. At the same time a 400µm low autofluorescence fiber was implanted into the injection site for fiber photometric assessment of ChAT+ neuron fiber activity across state changes. In addition, an LFP electrode was inserted into the frontal cortex and EMG electrodes attached to the nuchal muscle to allow polysomnography recording via telemetry transmitter (DSI, INC). After at least 1 month of recovery, the animals were connected to a laser emitter/photoreceptor apparatus and activity-dependent GCaMP6 fluorescence was recorded during multiple states. Low intensity light flashes (15-30µW) were used to prevent GCaMP bleaching. In addition, the photometry apparatus includes a 405 nm LED channel as control for movement artifacts. The two output signals were projected onto a photodetector (model 2151 femtowatt photoreceiver; Newport), and subsequently separated for analysis. Signals were collected at a sampling frequency of 381 Hz ^64^ for stimulus-evoked responses and at 0.1Hz for sleep/wake cycle monitoring. We expressed the 470nm GCaMP6 signal amplitude as a percent change from baseline activity during NREM sleep (dF = F at time t – mean NREM baseline F)/mean NREM baseline F) and then normalized the 470 nm signals to 405 nm control fluorescence by calculating (dF_470nm_−dF_405nm_) as previously described ^64,65^. Data were acquired using TDT software and analyzed in Spike2. Animals were tested in a chamber to which they had been extensively habituated or in their home cage during wake/sleep transitions, generally 1-2 hr sessions. For quantification of sleep-wake dependent changes in fluorescence, the correlation between LFP delta r.m.s. amplitude and fluorescence was calculated over at least 3 full state changes. In addition, dF relative to NREM basal activity was calculated. At the end of the experiment, brains were double IHC labeled for ChAT and GFP to confirm specificity of infection to ACh neurons.

### Pharmacology

Finally, as an initial step toward understanding mechanism of state-dependent synaptic change, a subset of the animals used in other manipulations (LEC: n=6, OB: n=3) were injected systemically with scopolamine (0, or 0.5mg/kg; within animal design) and OB or LEC evoked potentials and state changes monitored for 2-3 hours post-injection as described above.

### Experimental Design and Statistical Analysis

Experiment 1 involved n=6 LEC AAV-ChR infected rats and n=4 OB AAV-ChR infected chronically recorded rats and n=5 OB AAV-ChR infected urethane anesthetized rats. The design was a within animal state-dependent assay, combined with a between animal comparison of pathway specific effects. The primary within animal measure was a correlation of EP slope versus LFP delta r.m.s. amplitude. Correlations in each animal were converted to z-scores for comparison across groups with ANOVA and post-hoc tests. A secondary measure was a within animal comparison of LFP beta band coherence between LFP’s recorded in the PCX and LEC in a subset of the animals in the main experiment (n=4). A paired t-test was used to compare state-dependent changes in PCX-LEC coherence.

Experiment 2 involved analysis of state-dependent changes in the activity of basal forebrain ChAT+ neurons using fiberphotometry and GCaMP6 fluorescence (n=3). The primary within animal measure was a correlation of GCaMP6 fluorescence versus LFP delta r.m.s. amplitude. The significance of r was determined for each individual animal. In addition, the mean correlation across animal was calculated using z-score converted r’s for comparison against a mean of zero with a t-test.

Experiment 3 involved analysis of the effects of scopolamine on the state-dependent modulation of EP slope (LEC: n=6; OB: n=3). The primary within animal measure was a correlation of EP slope versus LFP delta r.m.s. amplitude. Correlations in each animal were converted to z-scores for comparison between drug and no drug with a paired t-test. Effects of drug on EP slope alone within state were compared with a paired t-test.

In all cases, results were considered significant if the probability of the statistical test outcome was < 0.05.

The datasets generated during the current study are available from the corresponding author on reasonable request.

## Acknowledgements

The authors thank Emmanuelle Courtiol and Brett East for assistance with viral transfections and E.C. for comments on the manuscript. This work was funded by R01-DC003906 to D.A.W.

## Author Contributions

D.A.W., M.J., M.I., A.P. and C.T. collected the data. D.A.W., M.J. and C.T. analyzed the data. D.A.W. prepared the figures. D.A.W. and C.T. wrote the manuscript.

